# Multimodal private information and reward value shape information use during ant foraging

**DOI:** 10.64898/2026.11.10.765227

**Authors:** Yu Tazuhara, Yuka Ito, Masaru K. Hojo

## Abstract

Animals use both private information acquired through experience and social information produced by conspecifics when making behavioural decisions. The combination of sensory cues and their associated reward value may influence how animals weigh private information against social information. We investigated information use during foraging in the ant *Pristomyrmex punctatus* using Y-maze experiments with learned visual and olfactory cues and trail pheromones. Workers learned both types of private cue and showed no significant preference between them when they were placed in conflict. When private information conflicted with a high-concentration pheromone trail, workers preferred the pheromone-marked route following visual, olfactory or multimodal training. At a lower pheromone concentration, route choice depended on training modality: workers trained with either unimodal cue preferred the pheromone trail, whereas workers trained with multimodal cues (combined visual and olfactory cues) showed no significant preference between the routes. When learned visual and olfactory cues were placed in conflict, workers favoured the cue paired with the higher sucrose concentration under both reciprocal reward assignments. In contrast, workers still preferred a pheromone trail when it had been associated with a lower-quality reward than the competing multimodal cues. These results show that ants adjust their use of learned private information according to expected reward and suggest that multimodal private information can influence the balance between private and social information during foraging.

**Summary statement:** Ants adjust their use of learned cues according to reward value, while multimodal cues can influence choices when private information conflicts with pheromone trails.

## INTRODUCTION

Animals make behavioural decisions in environments where multiple sources of information are simultaneously available. These sources often originate from different sensory modalities, such as visual, olfactory, or tactile cues, and animals frequently integrate them to guide behaviour. Multimodal information can improve the reliability of environmental assessment because different cues may provide complementary or redundant information about the same target (Partan and Marler, 1999; Hebets and Papaj, 2005). Multimodal cues can improve decision accuracy in foraging and reduce uncertainty in navigation (Kulahci et al., 2008; Nardini et al., 2008).

Information used for decision-making can also be classified according to its origin. Private information is acquired through an individual’s own experience, such as learned associations between environmental cues and resource locations, whereas social information is obtained from other individuals, for example through signals or behavioural cues produced by conspecifics (Danchin et al., 2004). Private information can provide direct and potentially reliable information about current conditions, but acquiring it requires personal sampling and may entail costs in time, energy or risk. Social information can reduce these sampling costs, but its reliability depends on how accurately it reflects current environmental conditions (Grüter and Leadbeater, 2014). Animals therefore flexibly balance private and social information according to the reliability and costs of the available information. In particular, reliance on social information often increases when private information is uncertain (Laland, 2004; Kendal et al., 2005). This raises the possibility that multimodal private information may influence not only individual learning and decision making, but also the relative use of private and social information. If combining multiple sensory cues increases the reliability of private information, individuals may be less likely to rely on socially provided information when the two sources conflict. The sensory modalities underlying private information may therefore influence how individuals balance private and social information.

Social insects provide excellent systems for studying this question because their collective behaviour emerges from the interaction between individual experience and socially transmitted signals. In ants, individual foragers can learn environmental cues associated with food sources, including visual landmarks and odours, and use these cues as private information during subsequent foraging trips (Steck et al., 2011; Oberhauser et al., 2019). At the same time, successful foragers of some species deposit trail pheromones that provide social information about routes to food sources and recruit nestmates to them (Wilson, 1962; Czaczkes et al., 2015). Because both learned environmental cues and pheromone trails can guide route choice, experienced foragers may encounter situations in which private and social information indicate different options.

Ants can flexibly adjust the relative weight given to these information sources. In *Lasius niger*, experienced foragers generally prioritise learned route information over conflicting pheromone trails (Grüter et al., 2011). However, when visual conditions become less favourable and private navigational information becomes less reliable, ants increase their reliance on pheromone trails, consistent with a “copy-when-uncertain” strategy (Jones et al., 2019). The information content itself can also affect this weighting: private information may contain multiple dimensions, such as both resource location and quality, whereas pheromone trails can provide strong directional information but comparatively limited information about resource quality (Czaczkes et al., 2019). Thus, the relative use of private and social information depends not simply on their source, but also on the reliability and content of the information available.

Experimental studies of private–social information conflict have often isolated components of private information, such as route memory or individual sensory cues, to determine their relative weight. In natural environments, however, private information is often composed of multiple sensory cues available simultaneously, such as visual landmarks together with odours associated with food sources (Buehlmann et al., 2020b). Examining how multimodal private information is used when it conflicts with social information can help explain how ants weigh competing information sources during route choice.

In this study, we investigated information-use strategies during foraging in the ant *Pristomyrmex punctatus*. This species uses chemical trails for recruitment and frequently relocates its nests (Tsuji, 1988b; Janssen et al., 1997). Its trail pheromone has been identified as 6-pentyl-2-pyrone, which is secreted by the poison gland (Janssen et al., 1997). This makes *P. punctatus* a useful model for examining interactions between individual learning and pheromone communication. Using Y-maze experiments, we first examined whether workers learn visual and olfactory cues as private information and whether they preferentially use either modality. We then tested how learned visual, olfactory and combined visual–olfactory private information was weighted against trail pheromone information when the two sources conflicted. Finally, we manipulated reward quality to examine whether the relative use of these information sources changes with previous experience. By experimentally manipulating the sensory modalities and reward value associated with learned information, we aimed to determine how multimodal private information shapes the balance between private and social information during ant foraging.

## MATERIALS AND METHODS

### Study species and colony maintenance

Colonies of the ant *Pristomyrmex punctatus* were collected on the Kobe Sanda Campus of Kwansei Gakuin University (Hyogo, Japan). *Pristomyrmex punctatus* is a queenless ant in which workers reproduce thelytokously. Reproductive and non-reproductive tasks are associated with age polyethism: younger workers reproduce and perform intranidal tasks, whereas older workers cease reproduction and shift to extranidal tasks such as foraging (Tsuji, 1988a, 1990; Tsuji and Dobata, 2011). Each colony was housed in a plastic container (440 × 295 × 75 mm) whose inner walls were coated with Fluon to prevent escape. Several water-filled test tubes (16 × 100 mm), plugged with cotton and wrapped in aluminium foil to exclude light, were placed inside the container and served as artificial nests. Colonies were maintained in an incubator at 25°C under a 16 h:8 h light:dark cycle. Ants were provided with water ad libitum and fed an artificial diet three times per week with two protein-to-carbohydrate ratios (P:C = 1:3 and 3:1), based on the synthetic diet described by Dussutour and Simpson (2008). Prior to behavioural experiments, 40–50 foraging workers were transferred to a circular plastic container and deprived of food for four days to increase foraging motivation. Starved workers were then divided into two groups: a training group consisting of approximately 10 workers and a recipient group consisting of approximately 40 workers used for trophallaxis. After each training visit, ants were allowed to unload food to nestmates in the recipient group. The number of colonies and workers differed among experiments and treatment groups; detailed sample sizes are provided in each figure and its corresponding legend.

### Solutions, stimuli and pheromone preparation

A 1.0 M sucrose solution was used as the food reward in all experiments unless otherwise stated.

Olfactory stimuli. Paper overlays and sucrose solutions were scented with nonanal or 1-octanol. Paper overlays were impregnated with odour by placing them overnight in a sealed glass container together with a beaker containing 100 µl of the odorant. During training, sucrose solutions were also scented with 0.005% of the corresponding odour. Visual stimuli. Visual cues consisted of blue or green coloured paper placed at the end of the runway during training (Fig. 1). Blue (R 0, G 75, B 225) and green (R 0, G 225, B 75) stimuli were printed on paper sheets using a colour printer (SC-PX1V, Epson, Japan).

**Fig. 1.**
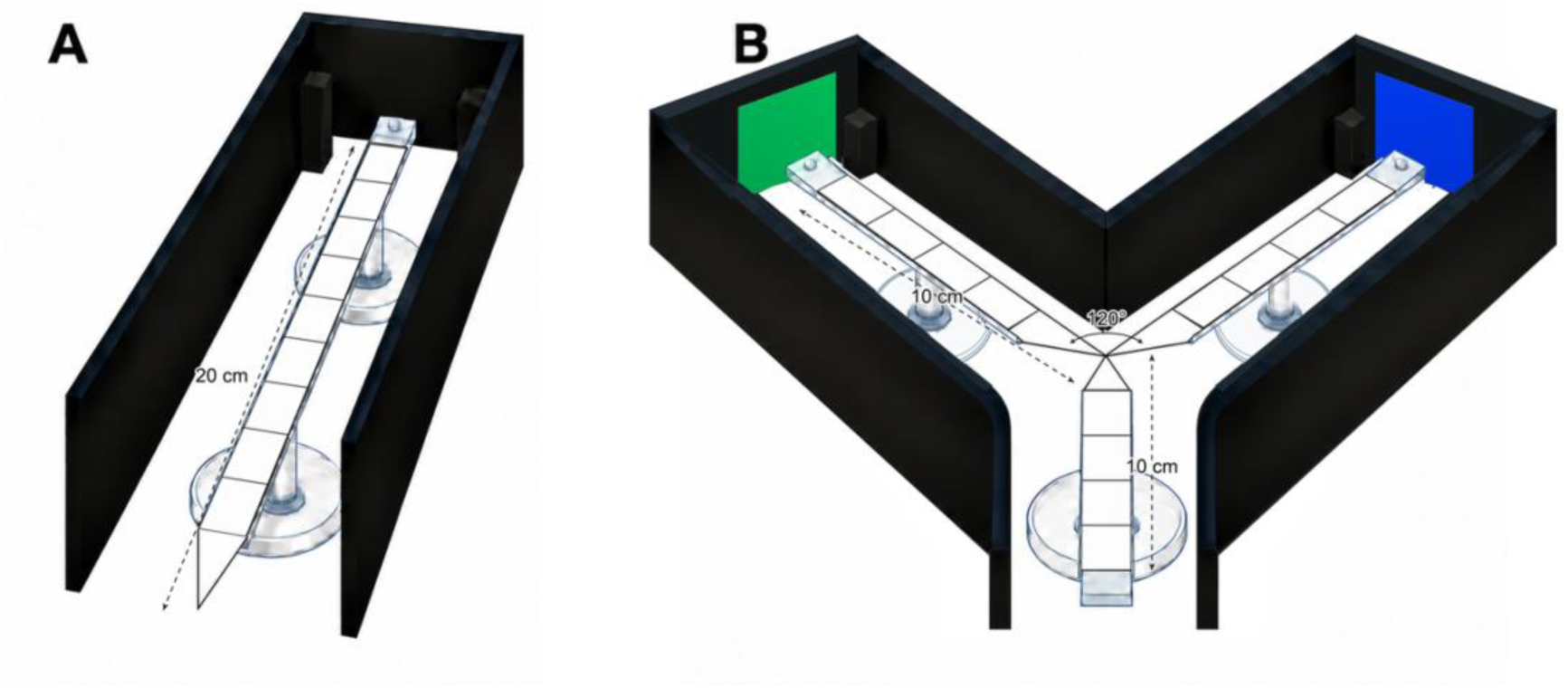
Experimental apparatus used for training and route-choice tests. Left: straight runway used during training. Workers entered from the colony and received a sucrose reward at the end of the 20 cm runway. Right: Y-maze used for route-choice tests. The stem and each arm were 10 cm long and 1 cm wide. Initial and final decisions were recorded at 2 and 8 cm from the bifurcation, respectively. Black walls surrounded the apparatus to minimise uncontrolled visual landmarks.

Trail pheromone. The trail pheromone of *P. punctatus*, 6-pentyl-2-pyrone (Tokyo Chemical Industry, Japan), was dissolved in *n*-hexane and stored at −30°C until use. For dose–response tests, pheromone solutions were prepared at 0.2, 0.4, 0.8, 2, 4, 8, 12 and 20 pg/µl; a solvent-only treatment served as the 0 pg/µl condition. During experiments, 10 µl of pheromone solution was applied along a 10 cm paper overlay using a microsyringe, while 10 µl of *n*-hexane served as the solvent control.

### Experimental setup

Most behavioural experiments consisted of a training phase followed by a test phase. Training was conducted on a straight runway (20 cm long, 1 cm wide) with a sucrose reward placed at the end, whereas testing was conducted on a Y-maze (stem and arms 10 cm long, 1 cm wide) following the design described by Czaczkes (2018). Both the straight runway and the Y-maze were constructed from acrylic plates covered with disposable paper overlays. To minimise the use of uncontrolled visual landmarks, the apparatus was surrounded by black walls (Fig. 1). In experiments involving runway training, ants that did not return to the starting point within 10 min were not tested further and were excluded from the analysis. In Y-maze tests, ants that did not make a choice within 5 min were excluded from the analysis. To quantify route-choice behaviour, two decision lines were defined on each arm of the Y-maze. Crossing a line 2 cm from the bifurcation was recorded as the initial decision, whereas crossing a line 8 cm from the bifurcation was recorded as the final decision.

### Experimental procedures

#### Concentration dependence of behavioural responses to trail pheromone

To determine the pheromone concentrations used in subsequent experiments, we examined workers’ behavioural responses to different concentrations of trail pheromone in the Y-maze. In each test, one arm of the Y-maze was assigned as the pheromone arm and the other as the control arm. Ten microlitres of pheromone solution were applied to the pheromone arm using a microsyringe, whereas 10 µl of *n*-hexane was applied to the control arm. The side receiving pheromone was randomised between trials. After pheromone application, the Y-maze was connected to the starved colony using a bridge. When a worker entered the maze, the bridge was removed, and the ant’s route choice was recorded.

### Olfactory learning

Workers were trained to associate an odour cue with a sucrose reward. During training, the straight runway was covered with paper scented with either nonanal or 1-octanol. Individual ants were allowed to enter the runway via a bridge. Once an ant reached the reward, it was marked on the abdomen for identification using a paint marker (PX-20; Mitsubishi, Japan). After returning to the starting position, the ant was transferred to the recipient group and allowed to unload food for 3 min. Each ant completed four training visits. Memory tests were conducted 5 min and 60 min after the final training visit using the same individual. Between the two tests, ants were returned to the recipient group. In each test, ants were allowed to choose between a scented arm and an unscented arm of the Y-maze, with the position of the scented arm randomised between trials. To test for innate odour preferences, additional Y-maze tests were conducted without prior training, in which ants chose either between scented and unscented arms or between arms scented with nonanal and 1-octanol.

### Visual learning

Visual learning experiments followed the same general procedure as the olfactory learning experiments. During training, either a blue or green visual cue was presented at the end of the straight runway, and each ant completed four training visits in which the cue was associated with a sucrose reward. All workers were tested 5 min after the final training visit; in a subset of experimental replicates, the same workers were additionally retested at 60 min. In each test, ants were allowed to choose between the trained colour and the alternative colour in the Y-maze, with cue positions randomised between trials. To assess innate colour preferences, additional control tests were conducted without prior training. In these tests, ants chose either between a coloured arm and a black arm, or between blue and green arms.

### Cue conflict experiments

To examine how learned private information was weighted against other information sources, we conducted a series of cue-conflict experiments.

First, to compare two learned private cues, workers were trained separately with a visual cue and the olfactory cue nonanal. Each cue type was presented four times in random order, resulting in a total of eight training visits. In the subsequent Y-maze test, one arm contained the trained visual cue, whereas the other contained nonanal together with the alternative, untrained colour. Thus, each arm contained only one of the two learned cues.

Next, to examine conflict between unimodal private information and trail pheromone, workers were trained for four visits with either a visual cue alone or nonanal alone. During testing, one arm contained the trained private cue and received *n*-hexane as a solvent control, whereas the other arm received trail pheromone. In experiments using visual private information, the pheromone arm also contained the alternative, untrained colour. Pheromone solutions of 8 or 0.8 pg/µl were used.

Finally, to examine conflict between multimodal private information and trail pheromone, workers were trained for four visits with visual and olfactory cues presented simultaneously. During testing, one arm contained the trained multimodal cues (nonanal and the trained colour) together with *n*-hexane, whereas the other contained trail pheromone together with the alternative colour. Pheromone solutions of 8 or 0.8 pg/µl were used. All cue-conflict tests were conducted 5 and 60 min after the final training visit using the same individual.

### Reward manipulation experiment

To examine whether reward quality affected information use, we manipulated the sucrose concentration associated with different information sources during training.

For comparisons between visual and olfactory private information, workers received four training visits with each cue, presented in random order for a total of eight visits. In one condition, the visual cue was associated with 0.1 M sucrose and nonanal with 1.0 M sucrose. In the reciprocal condition, the visual cue was associated with 1.0 M sucrose and nonanal with 0.1 M sucrose. During testing, one arm contained nonanal together with the alternative, untrained colour, whereas the other arm contained the trained visual cue.

In a third experiment, workers were trained to associate multimodal private information (visual + olfactory cues) with 1.0 M sucrose and an 8 pg/µl pheromone trail with 0.1 M sucrose. The workers received four training visits with each cue, presented in random order for a total of eight visits. During testing, ants were allowed to choose between an arm containing the trained multimodal cues and an arm containing trail pheromone together with the alternative colour.

All reward-manipulation tests were conducted 5 and 60 min after the final training visit using the same individual.

### Statistical analysis

All analyses were conducted in R version 4.4.2 (R Core Team, 2024) using generalised linear mixed models (GLMMs) with a binomial error distribution and logit link implemented in the glmmTMB package (Brooks et al., 2017). The final arm choice in the Y-maze was used as the binary response variable. Although both initial and final choices were recorded, only 13 of the 1602 tests included in the switching summary (0.8%) involved a switch between the initial and final decision lines; therefore, only final choices were analysed.

Colony identity was included as a random intercept in all models. When the same worker contributed observations at both 5 and 60 min, worker identity nested within colony was additionally included as a random intercept. The side on which the focal cue was presented was included as an additional categorical fixed effect to account for possible side bias. Colour identity (blue or green) was included as an additional categorical fixed effect in cue-conflict and reward-manipulation experiments involving visual cues.

For the pheromone dose–response experiment, pheromone concentration was treated as a categorical fixed effect. For the olfactory and visual learning experiments, stimulus identity, time after training and their interaction were included as fixed effects. Innate-preference tests were analysed using stimulus identity and cue side as appropriate. For the direct comparison between learned visual and olfactory cues, time after training was included as the focal fixed effect. For the private-information versus pheromone experiments, training modality, time after training and their interaction were included as fixed effects.

For the reciprocal visual–olfactory reward-manipulation experiment, data from the two reward assignments were analysed jointly using visual-cue choice as the binary response, with reward assignment (higher reward associated with the visual or olfactory cue), time after training and their interaction included as fixed effects. The multimodal–pheromone reward experiment was analysed separately, using multimodal-cue choice as the binary response and time after training as the focal fixed effect.

Model fit and dispersion were assessed using simulated residuals generated with the DHARMa package version 0.4.7 (Hartig, 2024). The significance of fixed effects was evaluated using Type II Wald χ² tests implemented with the Anova function of the car package (Fox and Weisberg, 2019). Estimated marginal means were calculated using the emmeans package version 2.0.0 (Lenth and Piaskowski, 2025). Two-sided tests against a choice probability of 0.5 were performed on the logit scale, and estimated probabilities and 95% confidence intervals were back-transformed for presentation. Holm correction was applied across the nine pheromone concentrations, across the two stimuli in the corresponding innate-preference tests, and across the two test times within each stimulus, training modality or reward assignment, as applicable. For Fig. 6A, tests against 0.5 were obtained from the same joint model used to test the effect of reward assignment. Reported *P*-values for these multiple comparisons are adjusted values.

## RESULTS

### Behavioural responses to trail pheromone

Workers responded to the trail pheromone 6-pentyl-2-pyrone, but the response depended on pheromone concentration (Fig. 2; GLMM: χ² = 23.06, d.f. = 8, *P* = 0.003). Significant preferences for the pheromone-treated arm were detected at 0.8, 4 and 8 pg/µl, with the highest choice probability observed at 8 pg/µl (tests against 0.5: 0.8 pg/µl, *P* = 0.042; 4 pg/µl, *P* = 0.009; 8 pg/µl, *P* < 0.001). At concentrations above 8 pg/µl, the proportion of workers choosing the pheromone-treated arm declined and no longer differed significantly from random choice (*P* > 0.05). We therefore used 0.8 pg/µl, the lowest concentration eliciting a significant preference, and 8 pg/µl, which elicited the highest response, in subsequent experiments.

**Fig. 2.**
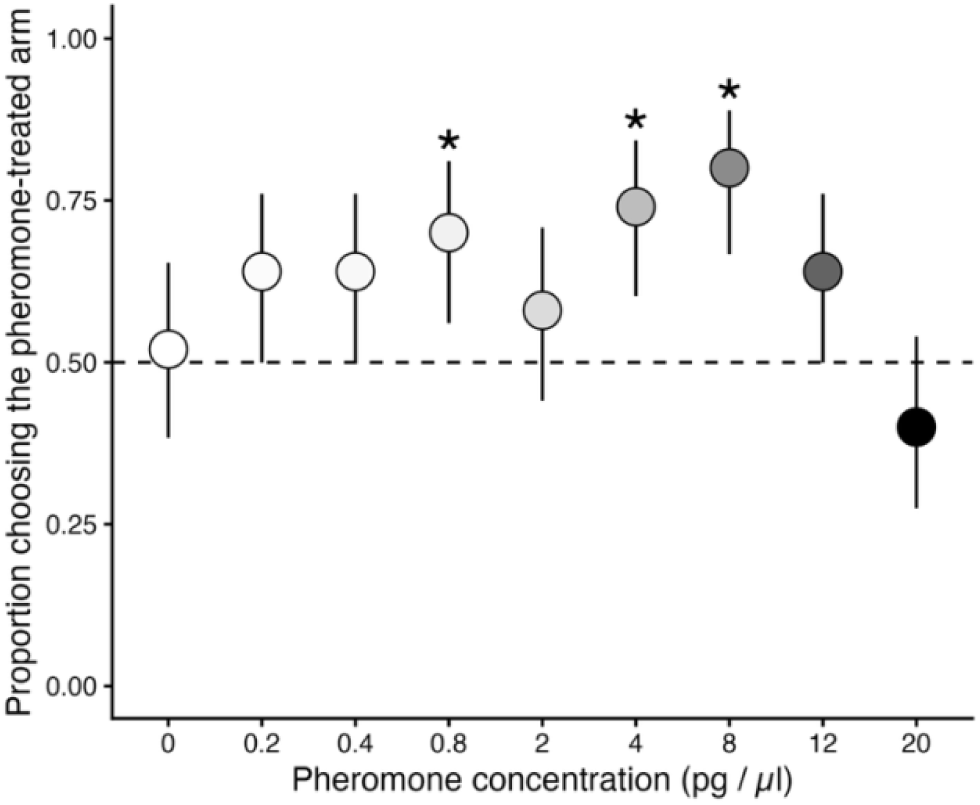
Behavioural responses of *Pristomyrmex punctatus* workers to different concentrations of trail pheromone. Points show estimated probabilities of choosing the pheromone-treated arm, with 95% confidence intervals. The dashed horizontal line indicates random choice (0.5). Asterisks indicate significant deviations from 0.5 after Holm adjustment (*P* < 0.05). Each concentration was tested with 50 workers from five colonies.

### Ants use both visual and olfactory cues as learned private information

Workers showed no innate preference for either nonanal or 1-octanol over an unscented arm (Fig. S1A; tests against 0.5, *P* > 0.05). When nonanal and 1-octanol were presented directly against each other without prior training, workers also showed no preference between the two odours (Fig. S1B; *P* = 0.864).

After four training visits, workers learned the association between an odour cue and the food reward. Choice probabilities did not differ significantly between odours or test times (GLMM; odour × time: χ² = 0.35, d.f. = 1, *P* = 0.552; time: χ² = 0.03, d.f. = 1, *P* = 0.866; odour: χ² = 2.50, d.f. = 1, *P* = 0.114). Workers significantly preferred nonanal after both 5 and 60 min (Fig. 3A; *P* < 0.001 at both times). For workers trained with 1-octanol, the estimated probability of choosing the scented arm was 0.78 at 5 min and 0.74 at 60 min, but neither estimate differed significantly from 0.5 (Fig. 3B, P = 0.054 and 0.056, respectively). Nonanal was therefore used as the olfactory cue in subsequent experiments.

**Fig. 3.**
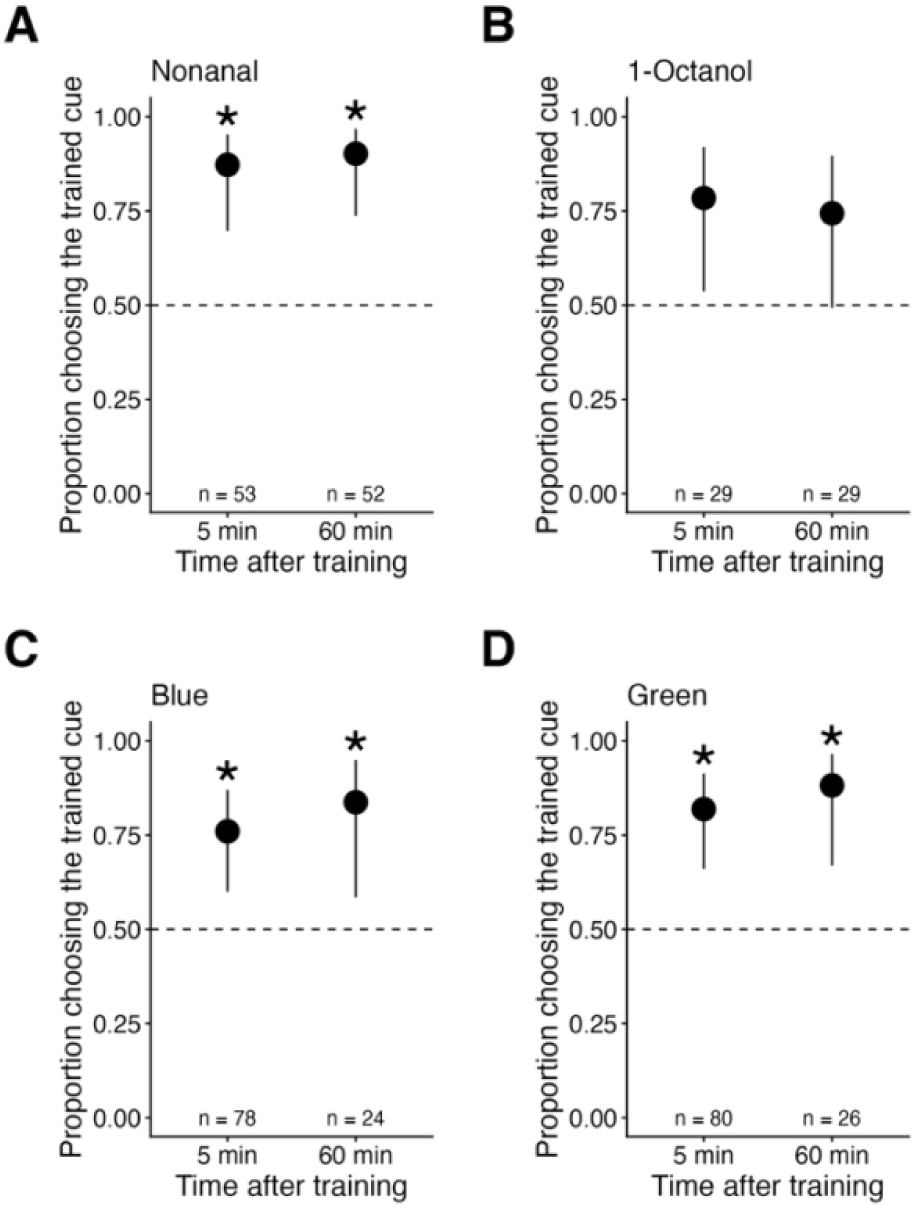
Learning and retention of olfactory and visual cues. Workers were trained to associate. (A) nonanal, (B) 1-octanol, (C) blue or (D) green cues with a sucrose reward and were tested 5 min after training. The same workers were retested at 60 min, except in the visual-learning experiment, in which only a subset was retested. Points show estimated probabilities of choosing the trained cue, with 95% confidence intervals. Dashed lines indicate random choice (0.5). Asterisks indicate significant deviations from 0.5 (*P* < 0.05), with Holm correction across the two test times within each stimulus. Numbers indicate the numbers of workers included in the analysis at each time point; three to six colonies were used.

Workers showed an innate response to visual cues. When blue or green was presented against a visually neutral black arm, the estimated marginal probability of choosing the coloured arm was significantly greater than chance (Fig. S2A; *P* = 0.003), and the response did not differ between blue and green (GLMM: χ² = 0.04, d.f. = 1, *P* = 0.834). When the two colours were evaluated separately, however, neither deviation from chance was significant (both *P* = 0.052). When blue and green were presented directly against each other without prior training, workers showed no preference between the two colours (Fig. S2B; *P* = 0.823).

After training, workers learned the association between visual cues and food reward. Choice probabilities did not differ significantly between colours or test times (GLMM; colour × time: χ² < 0.001, d.f. = 1, *P* = 0.986; time: χ² = 1.15, d.f. = 1, *P* = 0.283; colour: χ² = 0.82, d.f. = 1, *P* = 0.364). Workers significantly preferred the trained colour at both 5 and 60 min for both blue and green cues (Fig. 3C,D; all *P* < 0.05). Thus, *P. punctatus* workers can learn both olfactory and visual cues and use them as private information during subsequent route choice.

### Training modality affects the balance between private and social information

We first tested whether workers showed a preference between two learned private cues. After four training visits with a visual cue and four with an olfactory cue, workers showed no significant preference between the learned visual and olfactory cues at either test time (Fig. 4; GLMM, time: χ² = 0.74, d.f. = 1, *P* = 0.391; tests against 0.5, *P* = 1.0 at both times).

**Fig. 4.**
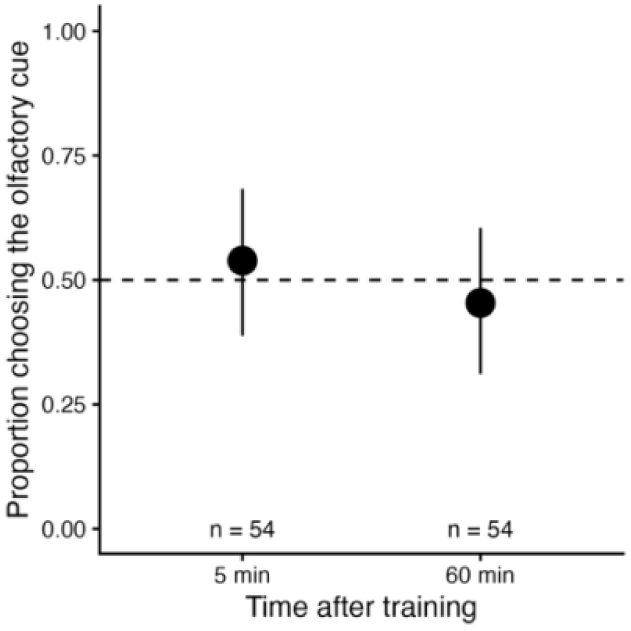
Relative use of learned visual and olfactory information. Workers were trained with visual and olfactory cues and subsequently given a choice between the two learned cues. Points show estimated probabilities of choosing the olfactory cue 5 and 60 min after training, with 95% confidence intervals. The dashed line indicates random choice (0.5). Numbers indicate the numbers of workers tested at each time point. Data were obtained from six colonies.

We next examined how training modality influenced decisions when learned private information conflicted with trail pheromone. At 8 pg/µl, workers significantly preferred the pheromone-treated arm following visual, olfactory and multimodal training at both test times (Fig. 5A; all *P* ≤ 0.0023). Choice probabilities did not differ significantly among training modalities or between test times (modality × time: χ² = 0.73, d.f. = 2, *P* = 0.694; time: χ² = 0.02, d.f. = 1, *P* = 0.893; modality: χ² = 1.05, d.f. = 2, *P* = 0.591).

**Fig. 5.**
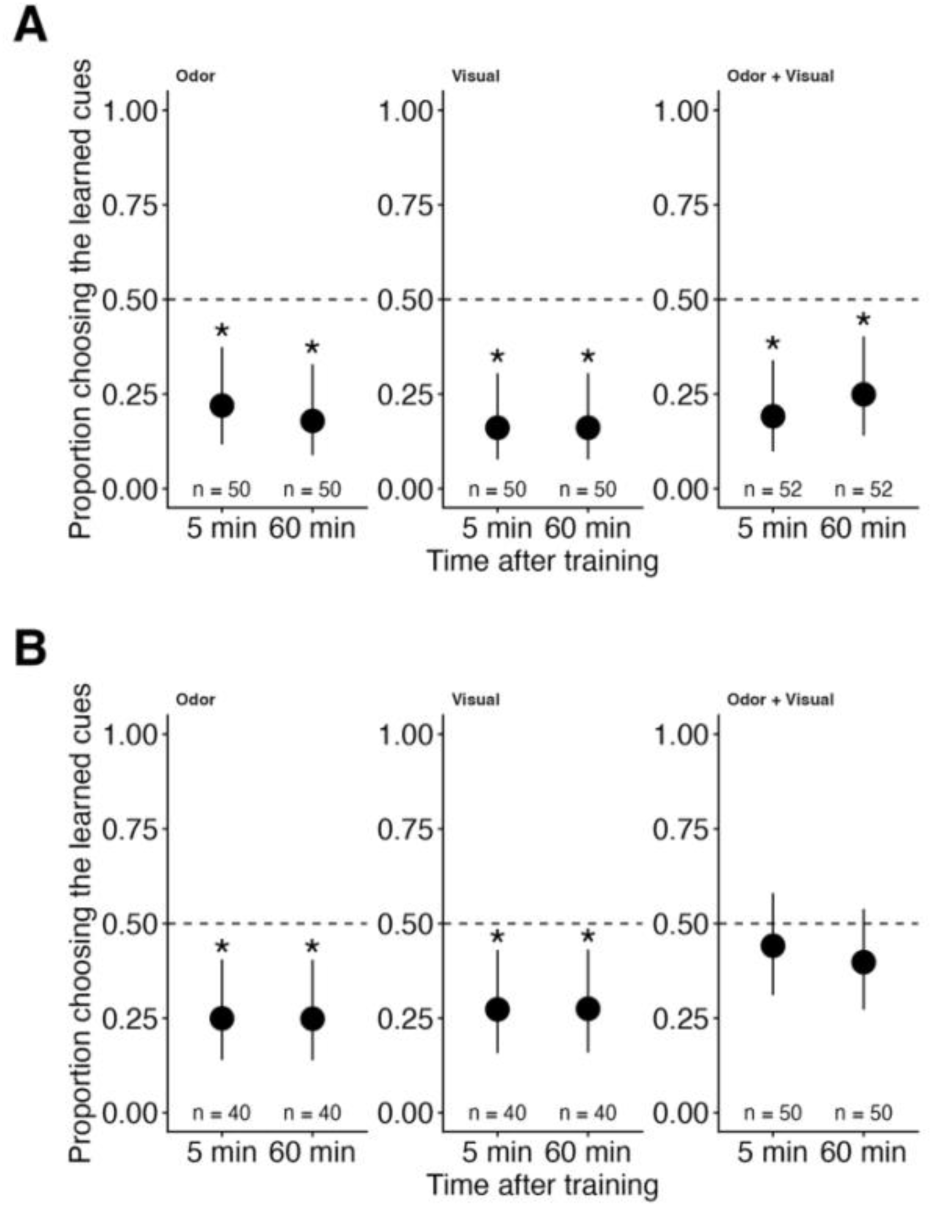
Use of unimodal and multimodal private information in conflict with trail pheromone. Workers were trained with an olfactory cue, a visual cue, or combined olfactory and visual cues and were subsequently tested against a pheromone-treated arm containing (A) 8 pg/µl or (B) 0.8 pg/µl 6-pentyl-2-pyrone. Points show estimated probabilities of choosing the learned private cue(s), with 95% confidence intervals. Values below 0.5 therefore indicate preference for the pheromone-treated arm. Asterisks indicate significant deviations from 0.5 (*P* < 0.05), with Holm correction across the two test times within each training modality. Numbers indicate the numbers of workers tested at each time point; four to six colonies were used.

At 0.8 pg/µl, however, choice probabilities differed significantly among training modalities (Fig. 5B; modality: χ² = 7.02, d.f. = 2, *P* = 0.030), whereas neither test time nor the modality × time interaction was significant (time: χ² = 0.08, d.f. = 1, *P* = 0.775; modality × time: χ² = 0.11, d.f. = 2, *P* = 0.947). Workers trained with visual or olfactory cues alone significantly preferred the pheromone-treated arm at both test times (visual: *P* = 0.012 at both times; olfactory: *P* = 0.005 at both times). In contrast, workers trained with combined visual and olfactory cues showed no significant preference between the learned multimodal cues and the pheromone-treated arm (5 min: *P* = 0.408; 60 min: *P* = 0.306).

### Flexible use of private information based on reward quality

Reward quality affected the relative use of visual and olfactory private information (Fig. 6A). In the joint model, reward assignment significantly affected visual-cue choice (χ² = 15.46, d.f. = 1, *P* < 0.001), whereas neither test time (χ² = 0.36, d.f. = 1, *P* = 0.550) nor the reward assignment × time interaction (χ² = 0.005, d.f. = 1, *P* = 0.944) was significant. When nonanal was associated with 1.0 M sucrose and the visual cue with 0.1 M sucrose, workers preferred the olfactory cue at both test times (tests against 0.5: 5 min, *P* = 0.0329; 60 min, *P* = 0.0305). When the reward assignments were reversed, workers preferred the visual cue (5 min, *P* = 0.0303; 60 min, *P* = 0.0395).

**Fig. 6.**
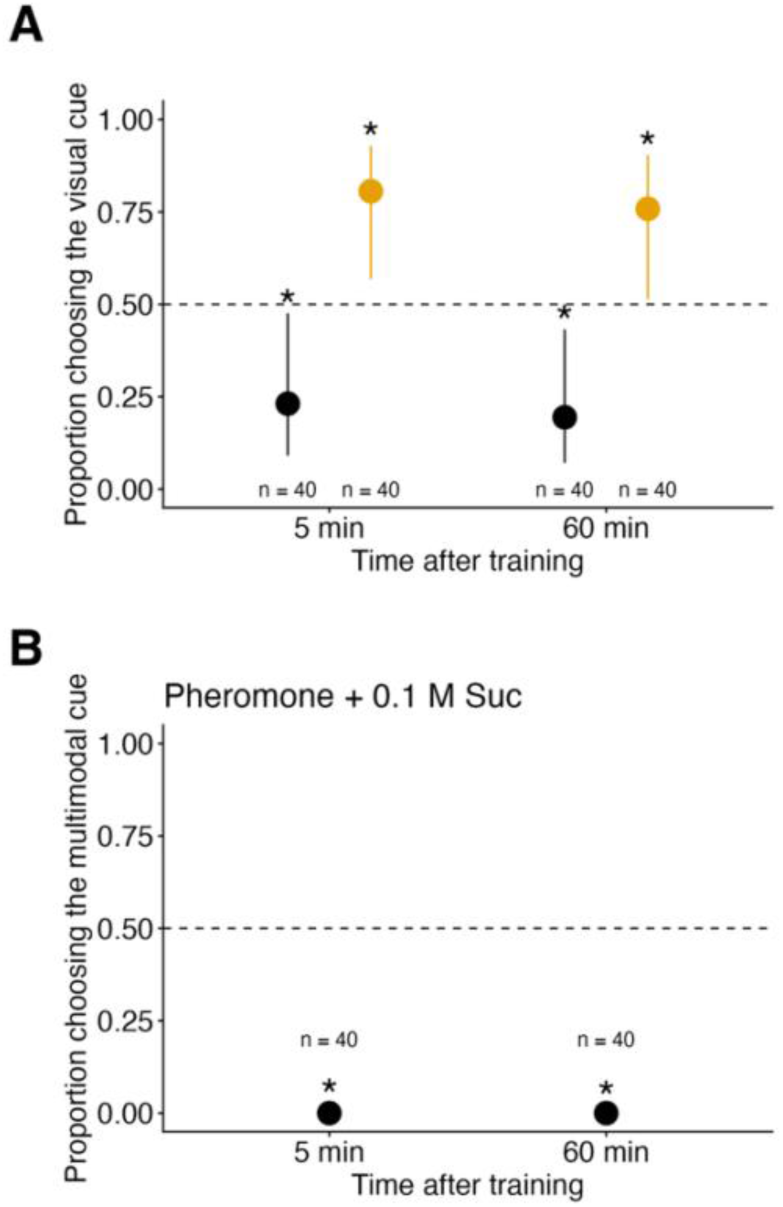
Effects of reward quality on subsequent information use. (A) Reciprocal visual–olfactory reward manipulation. Black points represent workers trained with the visual cue paired with 0.1 M sucrose and nonanal with 1.0 M sucrose; orange points represent the reverse assignment. Points show estimated probabilities of choosing the visual cue from a single model fitted jointly to both reward assignments. (B) Workers were trained with multimodal private cues paired with 1.0 M sucrose and an 8 pg/µl pheromone trail paired with 0.1 M sucrose. Points show estimated probabilities of choosing the multimodal cue. Error bars show 95% confidence intervals. The same workers were tested 5 and 60 min after training. Dashed lines indicate a choice probability of 0.5. Asterisks indicate deviations from 0.5 (*P* < 0.05), with Holm correction across the two test times within each reward assignment in (A) and across the two times in (B); they do not denote comparisons between reward assignments. Each plotted condition includes 40 workers per time point. In (A), the olfactory-high and visual-high treatments used five and four colonies, respectively; (B) used five colonies.

In contrast, when multimodal private cues were associated with 1.0 M sucrose and an 8 pg/µl pheromone trail with 0.1 M sucrose, workers strongly preferred the pheromone-treated arm at both test times (Fig. 6B; tests against 0.5: 5 min, *P* = 0.0025; 60 min, *P* = 0.0037). This preference did not differ significantly between test times (GLMM: χ² = 2.27, d.f. = 1, *P* = 0.132).

Together, these results show that workers adjusted their relative use of visual and olfactory private information according to expected reward quality, whereas a strong pheromone trail continued to dominate route choice even when it was associated with the lower-quality reward.

## DISCUSSION

In this study, we examined how *Pristomyrmex punctatus* workers use visual, olfactory and pheromone cues during route choice. Workers learned visual and olfactory cues, with preferences for the trained colours and nonanal persisting for at least 60 min. When private cues conflicted with trail pheromone, workers generally preferred the pheromone-marked route. At the lower pheromone concentration, route choice differed among training modalities: workers trained with unimodal cues preferred the trail, whereas those trained with combined visual and olfactory cues showed no significant route preference. Reward quality also affected choice between private cues, but workers still preferred the higher-concentration trail when it had been paired with a poorer reward than the competing multimodal cues. These results suggest that multimodality and learned reward value can influence information use, while a strong pheromone trail may remain preferred even when private cues predict a better reward.

### Reward value and multimodality in information use

The balance between private and social information may depend on both the reward associated with each source and the reliability of the information it provides. Here, reward value refers to the expected quality of food, whereas reliability refers to how consistently and accurately a cue indicates relevant environmental conditions, such as food location or travel direction. Reward value can guide route choice: *Lasius niger* follows odours associated with better food even when route memory indicates the opposite direction (Oberhauser et al., 2019), and information about better food can shift these ants from route memory to pheromone following (Czaczkes et al., 2019). Our reciprocal reward assignments show that *P. punctatus* can adjust its choice between private cues according to food quality. Yet workers still preferred the higher-concentration pheromone trail when it had been paired with poorer food. Under these conditions, attraction to the trail persisted despite the association between private cues and a better reward. This pattern suggests that a strong social signal can constrain the expression of reward-based preferences. Related limits have been reported in *L. niger*, which can learn to ignore, but not reliably avoid, trails paired with punishment (Wenig et al., 2021).

Increased reliability of private information offers a possible explanation for the effect of multimodality. Studies in insects and vertebrates show that combining sensory cues can improve navigational performance (Rossier et al., 2000; Steck et al., 2011; Buehlmann et al., 2020a; Jones and Moss, 2021; Yamada et al., 2021) and the accuracy of foraging decisions (Kulahci et al., 2008). In ants, *L. niger* relies more on pheromone trails under reduced illumination, consistent with increased social information use when private navigational information is uncertain (Jones et al., 2019). In our study, route choice differed among training modalities at 0.8 pg/µl, and workers trained with combined cues showed no significant trail preference. One interpretation is that visual and olfactory cues together made the rewarded cue combination easier to recognise, reducing uncertainty and reliance on the trail. Greater cue salience or stronger learned associations could also contribute to this pattern. Because cue reliability was not manipulated independently of modality, increased reliability remains a hypothesis for explaining the observed choices.

This interpretation also raises the question of what ants learn when visual and olfactory cues are presented together. Animals may learn one component more strongly than the other, or learn the combination as a distinct cue. Hawkmoths trained with a scented visual target later responded to the odour alone but showed little response to the visual component (Balkenius and Dacke, 2013). In desert ants, combined visual and olfactory landmarks accelerated learning; after repeated training with the combination, however, ants searched less accurately when either component was presented alone (Steck et al., 2011). These examples show that responses to multimodal cues need not reflect equal contributions from their components and may depend on how the combination is learned. In our study, workers showed no significant preference between visual and olfactory cues after learning them separately, but this does not establish how the cues were represented after joint training. We did not test individual components after multimodal training, so the observed choices could reflect associations with individual components, the combined cue, or both. Testing individual components and recombined cue pairings after joint training would help distinguish these possibilities.

### Why pheromone trails were generally prioritised

*Pristomyrmex punctatus* generally preferred trail pheromone over learned private cues in our experiments. This differs from the preference for route memory often reported in experienced *L. niger* foragers (Grüter et al., 2011), but the balance between private and social information varies among ant species. In a direct comparison, *L. niger* increasingly used learned environmental cues, whereas Argentine ants continued to rely mainly on chemical trails (Aron et al., 1993). Such variation may reflect differences in colony organisation and foraging ecology. Colony size is associated with recruitment strategy across ant species (Beckers et al., 1989), and species that frequently coordinate collective movement may benefit from strong responses to trails. *Pristomyrmex punctatus* colonies can comprise more than 10,000 individuals (Tsuji, 1995; Tsuji and Dobata, 2011), and chemical recruitment and frequent nest relocation are well documented (Tsuji, 1988b; Janssen et al., 1997). Coordinating these activities in large colonies may help explain the strong trail preference observed here. Comparisons among species under similar experimental conditions would help assess how ecology and previous experience shape information use.

The role of pheromone trails also depends on signal properties and behavioural context. In *L. niger*, pheromone and route memory together improve walking speed and straightness, suggesting that trails can help confirm a familiar route (Czaczkes et al., 2011). Trails can also convey negative information, as shown by the repellent ‘no entry’ signal in Pharaoh ants (Robinson et al., 2005), although avoidance of unsuitable food can arise from aversive memories, as reported in Argentine ants (Zanola et al., 2025). In our study, the estimated probability of choosing the pheromone-treated arm was lower at concentrations above 8 pg/µl and no longer differed significantly from chance. This pattern provides no evidence of active avoidance. Excessively strong stimulation or changes in orientation are possible explanations for the lower choice probabilities. Measuring naturally deposited trails and comparing responses during foraging and nest relocation would help establish the ecological relevance of the tested concentrations and clarify their effects on information use.

## Conclusion

Our findings suggest that multimodal private information can influence its use relative to social information during foraging. Workers adjusted their choices between private cues according to reward quality, while a strong pheromone trail remained preferred even when it had been associated with poorer food. Together, these results indicate that learned reward value and the properties of competing cues contribute to information use. The learning processes underlying multimodal cue use and the proposed contribution of reliability require direct tests. Reward effects on private–social choice also warrant further study because only one reward assignment was tested at the higher pheromone concentration. Testing cue components after joint training, and varying reliability and reward value independently, would clarify when private experience changes reliance on social information.

## Acknowledgements

We thank Y. Hamaya, T. Kikuoka and T. Morimoto for field sampling and technical support. We also thank K. Morishima and K. Kosuge for their valuable support during the research.

## Funding

This study was supported by the Japan Society for the Promotion of Science KAKENHI under grant numbers 22H02705 and 25K09144, and the Kinoshita Memorial Foundation.

## Data availability

The data and analysis code supporting this study will be made available as a Zenodo repository upon journal publication.

## Competing interests

The authors declare no competing or financial interests.

## Use of AI tools

ChatGPT and Codex (OpenAI) were used to improve author-generated texts and the preparation of the apparatus schematic in Fig. 1 from photographs of the experimental setup. The authors reviewed and verified the text and schematic.

## Notes

### Competing Interest Statement

The authors have declared no competing interest.

## References

Aron, S., Beckers, R., Deneubourg, J. L. and Pasteels, J. M. (1993). Memory and chemical communication in the orientation of two mass-recruiting ant species. Insectes Soc. 40, 369– 380. 10.1007/BF01253900

Balkenius, A. and Dacke, M. (2013). Learning of multi-modal stimuli in hawkmoths. PLoS ONE 8, e71137. 10.1371/journal.pone.0071137

Beckers, R., Goss, S., Deneubourg, J. L. and Pasteels, J. M. (1989). Colony size, communication and ant foraging strategy. Psyche 96, 239–256. 10.1155/1989/94279

Brooks, M. E., Kristensen, K., van Benthem, K. J., Magnusson, A., Berg, C. W., Nielsen, A., Skaug, H. J., Maechler, M. and Bolker, B. M. (2017). glmmTMB balances speed and flexibility among packages for zero-inflated generalized linear mixed modeling. R J. 9, 378–400. 10.32614/RJ-2017-066

Buehlmann, C., Aussel, A. and Graham, P. (2020a). Dynamic multimodal interactions in navigating wood ants: what do path details tell us about cue integration? J. Exp. Biol. 223, jeb221036. 10.1242/jeb.221036

Buehlmann, C., Mangan, M. and Graham, P. (2020b). Multimodal interactions in insect navigation. Anim. Cogn. 23, 1129–1141. 10.1007/s10071-020-01383-2

Czaczkes, T. J. (2018). Using T- and Y-mazes in myrmecology and elsewhere: a practical guide. Insectes Soc. 65, 213–224. 10.1007/s00040-018-0621-z

Czaczkes, T. J., Grüter, C., Jones, S. M. and Ratnieks, F. L. W. (2011). Synergy between social and private information increases foraging efficiency in ants. Biol. Lett. 7, 521–524. 10.1098/rsbl.2011.0067

Czaczkes, T. J., Grüter, C. and Ratnieks, F. L. W. (2015). Trail pheromones: an integrative view of their role in social insect colony organization. Annu. Rev. Entomol. 60, 581–599. 10.1146/annurev-ento-010814-020627

Czaczkes, T. J., Beckwith, J. J., Horsch, A.-L. and Hartig, F. (2019). The multi-dimensional nature of information drives prioritization of private over social information in ants. Proc. R. Soc. B 286, 20191136. 10.1098/rspb.2019.1136

Danchin, É., Giraldeau, L.-A., Valone, T. J. and Wagner, R. H. (2004). Public information: from nosy neighbors to cultural evolution. Science 305, 487–491. 10.1126/science.1098254

Dussutour, A. and Simpson, S. J. (2008). Description of a simple synthetic diet for studying nutritional responses in ants. Insectes Soc. 55, 329–333. 10.1007/s00040-008-1008-3

Fox, J. and Weisberg, S. (2019). An R Companion to Applied Regression. 3rd edn. Thousand Oaks, CA: Sage.

Grüter, C. and Leadbeater, E. (2014). Insights from insects about adaptive social information use. Trends Ecol. Evol. 29, 177–184. 10.1016/j.tree.2014.01.004

Grüter, C., Czaczkes, T. J. and Ratnieks, F. L. W. (2011). Decision making in ant foragers (*Lasius niger*) facing conflicting private and social information. Behav. Ecol. Sociobiol. 65, 141–148. 10.1007/s00265-010-1020-2

Hartig, F. (2024). DHARMa: Residual Diagnostics for Hierarchical (Multi-Level / Mixed) Regression Models. R package version 0.4.7. 10.32614/CRAN.package.DHARMa

Hebets, E. A. and Papaj, D. R. (2005). Complex signal function: developing a framework of testable hypotheses. Behav. Ecol. Sociobiol. 57, 197–214. 10.1007/s00265-004-0865-7

Janssen, E., Hölldobler, B., Kern, F., Bestmann, H. J. and Tsuji, K. (1997). Trail pheromone of myrmicine ant *Pristomyrmex pungens*. J. Chem. Ecol. 23, 1025–1034. 10.1023/B:JOEC.0000006386.14952.d7

Jones, S., Czaczkes, T. J., Gallager, A. J., Oberhauser, F. B., Gourlay, E. and Bacon, J. P. (2019). Copy when uncertain: lower light levels increase trail pheromone depositing and reliance on pheromone trails in ants. Anim. Behav. 156, 87–95. 10.1016/j.anbehav.2019.08.007

Jones, T. K. and Moss, C. F. (2021). Visual cues enhance obstacle avoidance in echolocating bats. J. Exp. Biol. 224, jeb241968. 10.1242/jeb.241968

Kendal, R. L., Coolen, I., van Bergen, Y. and Laland, K. N. (2005). Trade-offs in the adaptive use of social and asocial learning. Adv. Study Behav. 35, 333–379. 10.1016/S0065-3454(05)35008-X

Kulahci, I. G., Dornhaus, A. and Papaj, D. R. (2008). Multimodal signals enhance decision making in foraging bumble-bees. Proc. R. Soc. B 275, 797–802. 10.1098/rspb.2007.1176

Laland, K. N. (2004). Social learning strategies. Learn. Behav. 32, 4–14. 10.3758/BF03196002

Lenth, R. V. and Piaskowski, J. (2025). emmeans: Estimated Marginal Means, aka Least-Squares Means. R package version 2.0.0. 10.32614/CRAN.package.emmeans

Nardini, M., Jones, P., Bedford, R. and Braddick, O. (2008). Development of cue integration in human navigation. Curr. Biol. 18, 689–693. 10.1016/j.cub.2008.04.021

Oberhauser, F. B., Schlemm, A., Wendt, S. and Czaczkes, T. J. (2019). Private information conflict: *Lasius niger* ants prefer olfactory cues to route memory. Anim. Cogn. 22, 355–364. 10.1007/s10071-019-01248-3

Partan, S. and Marler, P. (1999). Communication goes multimodal. Science 283, 1272–1273. 10.1126/science.283.5406.1272

R Core Team (2024). R: A Language and Environment for Statistical Computing. Version 4.4.2. Vienna: R Foundation for Statistical Computing. https://www.R-project.org/

Robinson, E. J. H., Jackson, D. E., Holcombe, M. and Ratnieks, F. L. W. (2005). Insect communication: “no entry” signal in ant foraging. Nature 438, 442. 10.1038/438442a

Rossier, J., Haeberli, C. and Schenk, F. (2000). Auditory cues support place navigation in rats when associated with a visual cue. Behav. Brain Res. 117, 209–214. 10.1016/S0166-4328(00)00293-X

Steck, K., Hansson, B. S. and Knaden, M. (2011). Desert ants benefit from combining visual and olfactory landmarks. J. Exp. Biol. 214, 1307–1312. 10.1242/jeb.053579

Tsuji, K. (1988a). Obligate parthenogenesis and reproductive division of labor in the Japanese queenless ant *Pristomyrmex pungens*: comparison of intranidal and extranidal workers. Behav. Ecol. Sociobiol. 23, 247–255. 10.1007/BF00302947

Tsuji, K. (1988b). Nest relocations in the Japanese queenless ant *Pristomyrmex pungens* Mayr. (Hymenoptera: Formicidae). Insectes Soc. 35, 321–340. 10.1007/BF02225809

Tsuji, K. (1990). Reproductive division of labour related to age in the Japanese queenless ant, *Pristomyrmex pungens*. Anim. Behav. 39, 843–849. 10.1016/S0003-3472(05)80948-0

Tsuji, K. (1995). Reproductive conflicts and levels of selection in the ant *Pristomyrmex pungens*: contextual analysis and partitioning of covariance. Am. Nat. 146, 586–607. 10.1086/285816

Tsuji, K. and Dobata, S. (2011). Social cancer and the biology of the clonal ant *Pristomyrmex punctatus* (Hymenoptera: Formicidae). Myrmecol. News 15, 91–99. https://myrmecologicalnews.org/cms/index.php?filename=volume15/mn15_91-99_printable.pdf&format=raw&option=com_download&view=download

Wenig, K., Bach, R. and Czaczkes, T. J. (2021). Hard limits to cognitive flexibility: ants can learn to ignore but not avoid pheromone trails. J. Exp. Biol. 224, jeb242454. 10.1242/jeb.242454

Wilson, E. O. (1962). Chemical communication among workers of the fire ant *Solenopsis saevissima* (Fr. Smith). 1. The organization of mass-foraging. Anim. Behav. 10, 134–147. 10.1016/0003-3472(62)90141-0

Yamada, M., Ohashi, H., Hosoda, K., Kurabayashi, D. and Shigaki, S. (2021). Multisensory-motor integration in olfactory navigation of silkmoth, *Bombyx mori*, using virtual reality system. eLife 10, e72001. 10.7554/eLife.72001

Zanola, D., Czaczkes, T. J. and Josens, R. (2025). Toxic bait abandonment by an invasive ant is driven by aversive memories. *Commun*. Biol. 8, 486. 10.1038/s42003-025-07818-1

