## Supplemental Figures for "Multimodal private information and reward value shape information use during ant foraging"

**Supplementary Information**

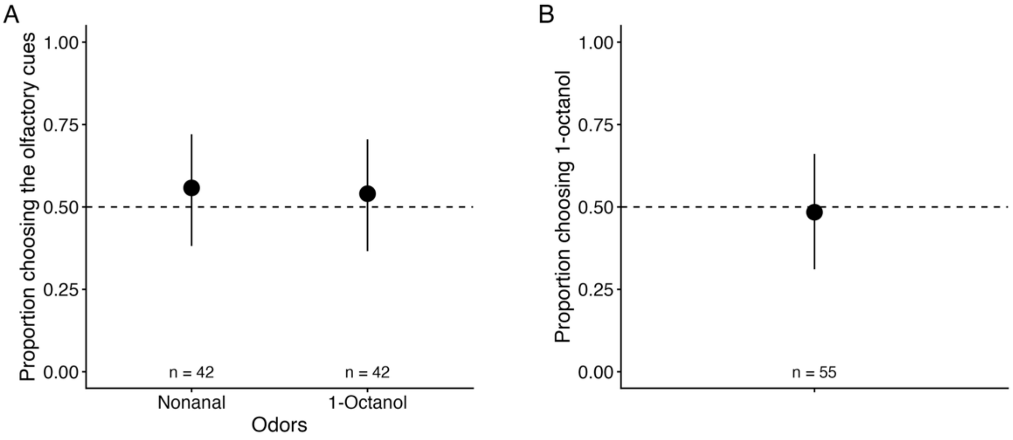


**Fig. S1. Innate responses to the olfactory stimuli used in the learning experiments.** (A) Choice between each odour and an unscented control without prior training. (B) Choice between nonanal and 1-octanol without prior training. Points show estimated choice probabilities with 95% confidence intervals; dashed lines indicate random choice (0.5). Numbers indicate the numbers of workers tested; three to six colonies were used.

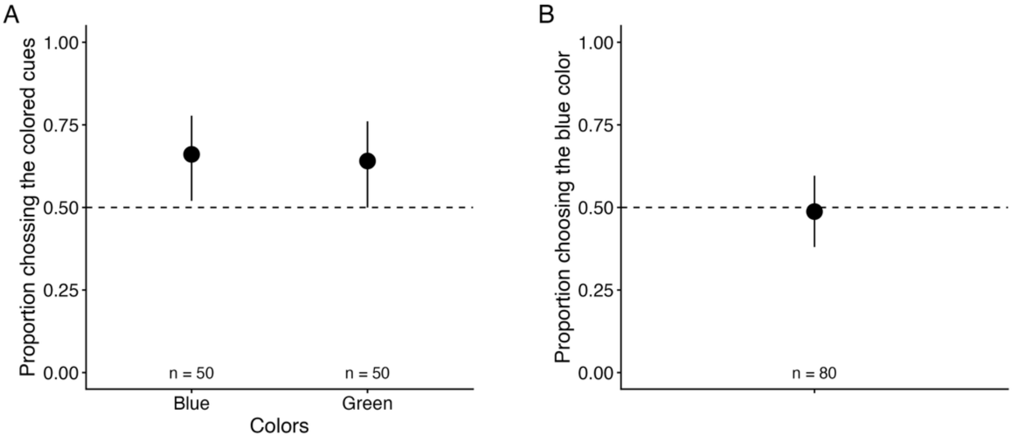


**Fig. S2. Innate responses to visual cues.** (A) Choice between a coloured arm (blue or green) and a black arm without prior training. (B) Choice between blue and green arms without prior training. Points show estimated choice probabilities with 95% confidence intervals. Dashed lines indicate random choice (0.5). In (A), workers showed an overall preference for the coloured arm, with no significant difference between blue and green. Numbers indicate the numbers of workers tested; four to five colonies were used.
